# Mouse Homolog of Human IRF8^G388S^ Mutation Provides Novel Insight into Osteoclastogenesis and Tooth Root Resorption

**DOI:** 10.1101/2023.04.23.537931

**Authors:** Amitabh Das, Sathish Kumar Yesupatham, Devon Allison, Himanshi Tanwar, JebaMercy Gnanasekaran, Bernice Kear, Xiaobei Wang, Sheng Wang, Christina Zachariadou, Yasaman Abbasi, Man-Kyo Chung, Keiko Ozato, Chengyu Liu, Brian L. Foster, Vivek Thumbigere-Math

## Abstract

Previously, we reported a novel mutation in the Interferon Regulatory Factor 8 (*IRF8*) gene associated with multiple idiopathic cervical root resorption (MICRR), an aggressive form of tooth root resorption mediated by increased osteoclast activity. The IRF8 G388S variant in the highly conserved C-terminal motif is predicted to alter the protein structure, likely impairing IRF8 function. To investigate the molecular basis of MICRR and IRF8 function, we generated *Irf8* knock-in (KI) mice using CRISPR/Cas9 modeling the human *IRF8^G388S^* mutation. The heterozygous (Het) and homozygous (Homo) *Irf8 KI* mice showed no gross morphological defects, and the development of hematopoietic cells was unaffected and similar to that in wild-type (WT) mice. The *Irf8 KI* Het and Homo mice showed no difference in macrophage gene signatures important for antimicrobial defenses and inflammatory cytokine production. Consistent with the phenotype observed in MICRR patients, *Irf8 KI* Het and Homo mice demonstrated significantly increased osteoclast formation and resorption activity *in vivo* and *in vitro* when compared to WT mice. The oral ligature inserted *Irf8 KI* Het and Homo mice displayed increased osteoclast-mediated alveolar bone loss and tooth root resorption compared to WT mice. The increased osteoclastogenesis noted in KI mice is due to the inability of mutant *Irf8* G388S isoform to negatively inhibit NFATc1-dependent transcriptional activation and downstream osteoclast specific transcripts. This translational study delineates the IRF8 domain important for osteoclast function and provides novel insights into the *IRF8* mutation associated with MICRR. *Irf8^G388S^* mutation mainly affects osteoclastogenesis while sparing immune cell development and function. The *Irf8 KI* mice serve as a novel translational model for studying the etiopathology of MICRR and developing targeted therapies for MICRR and other skeletal disorders mediated by increased osteoclast activity.

## Introduction

Tooth root resorption mediated by osteoclasts/odontoclasts is a normal physiologic process required for exfoliation of primary teeth to make space for the eruption of permanent teeth(^1^). However, root resorption of permanent teeth is largely pathological(^2–5^). Multiple Idiopathic Cervical Root Resorption (MICRR) is an aggressive form of pathologic root resorption that affects multiple teeth within the permanent dentition(^6–14^). MICRR lesions are often asymptomatic, non-carious, and lack overt gingival inflammation, deeper pocket depth, or tooth mobility that are associated with classical cases of periodontitis(^6–14^). MICRR lesions are often detected as incidental findings during routine dental examination and are resistant to interventions, resulting in tooth loss(^6,11,12^). To date, the etiology of MICCR remains largely speculative(^15^).

Previously, by examining a rare cohort of family members afflicted with MICRR, we identified a novel heterozygous IRF8 (G388S) mutation associated with increased osteoclast activity and susceptibility to MICRR(^16^). IRF8 is a member of the IRF family of transcription factors and plays an important role in myeloid cell differentiation, immune response, and transcription of type I IFN and IFN-inducible genes(^17–23^). Additionally, IRF8 functions as a negative regulator of osteoclast by inhibiting NFATc1, a master regulator of osteoclastogenesis(^16,23–25^). The identified C-terminal mutation within exon 9 of *IRF8* gene is immediately adjacent to the IRF association domain (IAD), a critical regulatory domain that mediates homo- and heterodimeric interactions among IRFs, and between IRFs and other transcriptional co-modulators(^26–29^). The IRF8 G388 residue is highly conserved among mammalian species(^16^). Conformational and bioinformatics analyses indicated that the G388S substitution could alter overall protein structure, likely impairing IRF8 heterodimerization with other transcription factors, including ETS, IRF and NFAT family members, ultimately affecting IRF8 function(^16^).

To establish the molecular basis of increased osteoclastogenesis and MICRR, we generated *Irf8* knock-in (KI) mice modeling the human *IRF8^G388S^* mutation. Consistent with the human findings and bioinformatics prediction, the *Irf8 KI* heterozygous (Het) and homozygous (Homo) mice recapitulated increased osteoclast formation and root resorption activity when compared to the wild-type (WT) mice. This comparative translational study delineates the IRF8 domain important for osteoclast function and provides novel insights into the *Irf8* mutation associated with MICRR. The *Irf8 KI* mouse model presents a valuable translational tool for further investigations into the etiology and potential therapies for MICRR.

## MATERIALS AND METHODS

### GENERATION OF *Irf8* Knock-in (KI) Mice

The Irf8 G388S KI mouse model was generated using CRISPR/Cas9 technology(^30^) on a C57BL/6 background. Briefly, a sgRNA (GCGGGCAAGAGCTGCGGTGC) was designed and synthesized using ThermoFisher’s Custom In Vitro Transcription Service. A single-strand donor oligonucleotides containing the desired mutation (GCAGGTAGAGCAGCTGTATGCCAGGCAGCTGGTGGAGGAAGCGGGCAAGAGCTGC GGTGC**<u>CTCT</u>**TCCCTGATGCCAGCCCTGGAGGAGCCCCAGCCGGACCAGGCTTTCCGCATGTTTCCGGAT; the four bold and underlined nucleotides were changed from the wild-type sequence for introducing the true and silent mutations) was ordered from IDT (https://www.idtdna.com/pages). The sgRNA (20 ng/ul) and donor oligonucleotides (100 ng/ul) were co-microinjected with Cas9 mRNA (20ng/ul, purchased from Trilink Biotechnologies) into the cytoplasm of zygotes collected from C57BL/6N mice (Charles River Laboratory). Injected embryos were cultured in M16 medium (MilliporeSigma) overnight in a 37° C incubator with 6% CO_2_. The next morning, embryos that had reached 2-cell stage of development were implanted into the oviducts of pseudopregnant surrogate mothers. The offspring’s born to the foster mothers were screened by PCR genotyping and Sanger Sequencing, and founder mice were established. The founder mice were subsequently bred with WT C57BL/6 mice for at least two generations to dilute any potential off-target mutations, and then used for the current study. Heterozygous mice with the same modification were mated to generate homozygous offspring.

All mice were maintained under specific pathogen-free conditions and all experiments were carried out in strict accordance with the recommendations outlined in the Guide for the Care and Use of Laboratory Animals from the National Institutes of Health (NIH). The protocol was approved by the Institutional Animal Care and Use Committee (IACUC) of the University of Maryland Baltimore School of Medicine (Protocol Numbers: 0218015 and 0121009; NHLBI H-0125) and this work adheres to the Animals in Research: Reporting In Vivo Experiments (ARRIVE) guidelines. Animals aged between 8-12 weeks were used for all experiments as noted in the figure legends. When possible, littermate controls were utilized for experiments. Both male and female mice and cells obtained from both genders were evaluated in this study. Mice were euthanized by CO_2_ inhalation. Mice were genotyped by standard PCR protocols.

### FLOW CYTOMETRY ANALYSIS

Flow cytometry was performed as previously described(^25,31^). Briefly, mice were sacrificed and whole blood samples were collected via cardiac puncture. After perfusion, the spleen was macerated in Hank’s balanced salt solution (HBSS) and filtered through a 70-μm cell strainer to obtain single cell suspensions. Bone marrow (BM) cells were harvested by flushing femurs and tibias. Following hypotonic lysis with ACK Lysing buffer, single cell suspensions were pre-incubated with Mouse BD Fc Block™ (clone 2.4G2), then stained with a cocktail of fluorescently labeled antibodies (Abs) (see **Table 1** for antibody list) for 30 minutes at 4**°**C. All antibodies were titrated for optimal dilution. After staining, the cells were washed with 1x PBS and then viability dye 7-AAD was used to exclude dead cells from analysis. Flow cytometry was performed on a 3 laser (488 nm, 407 nm, 640 nm) Cytek Aurora spectral cytometer (Cytek Biosciences), and the spectral data were unmixed based on single color beads controls using SpectroFlo software, then analyzed using SpectroFlo software. ‘‘Fluorescence minus one’’ controls were used when necessary to confirm the correct gates.

**TABLE 1:**
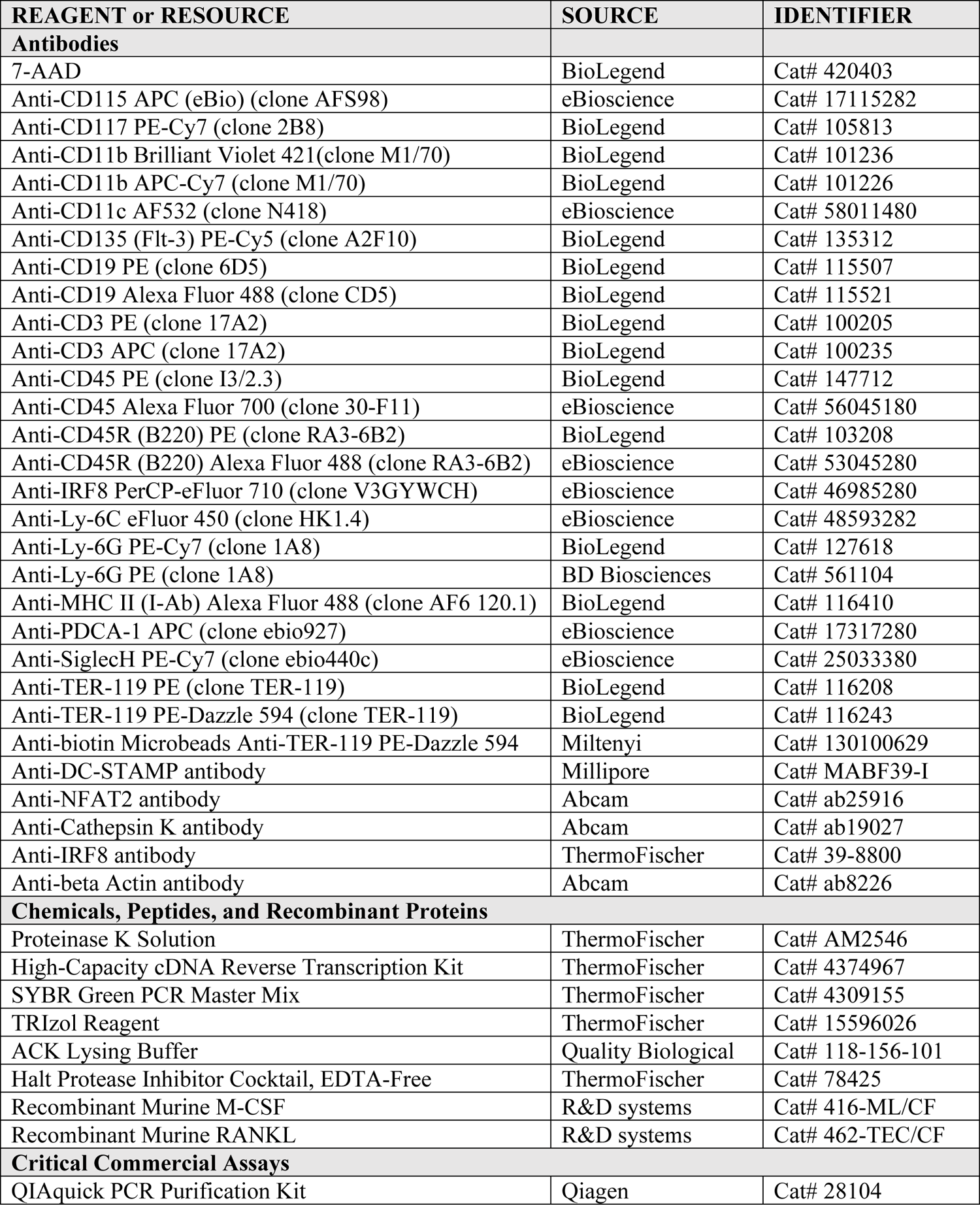
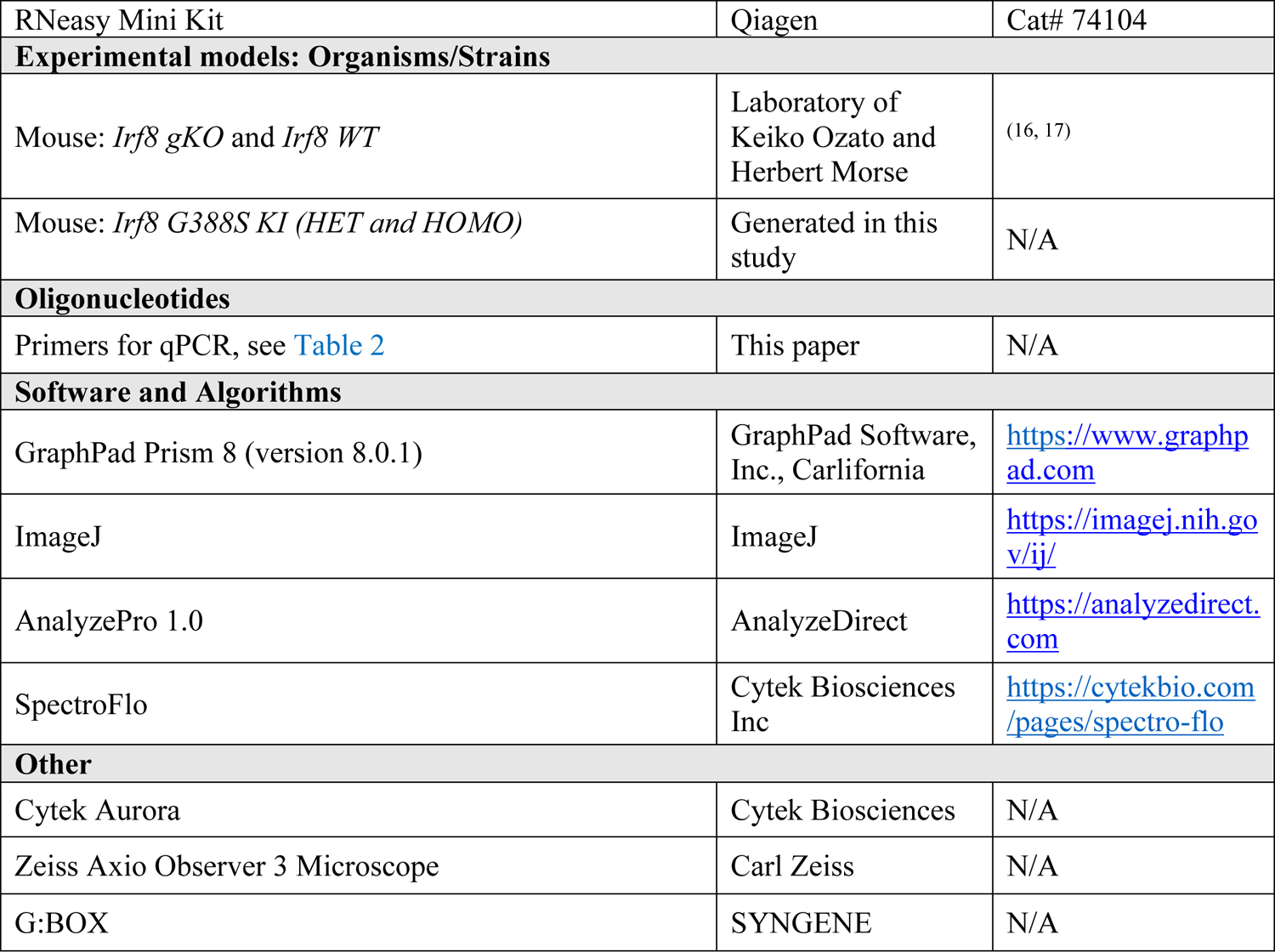
KEY RESOURCES

### *IN VITRO* OSTEOCLAST ASSAY

Osteoclast assays were performed as described previously(^16,25^). Briefly, primary BM cells were harvested from femurs and tibias of 8-12-week-old animals. After lysis of red blood cells using ACK Lysing Buffer (ThermoFisher Scientific), BM cells were cultured overnight in 100 mm tissue culture dishes (Corning) in osteoclast media (phenol red-free alpha-MEM (Gibco)) with 10% heat-inactivated fetal bovine serum (Hyclone), 25 units/mL penicillin/streptomycin (Invitrogen), 400 mM L-Glutamine (Invitrogen), and supplemented with 20 ng/ml M-CSF. The non-adherent cell population was recovered and re-plated in 12-well plates (Corning) at 5×10^5^ cells/cm^2^ and 6-well plates (Corning) at 1×10^6^ cells/ cm^2^ in osteoclast media supplemented with 20 ng/ml M-CSF. The following day, cells were treated with 40 ng/mL RANKL (R&D Systems). Subsequently, media were replaced every 2 two days with 20 ng/ml M-CSF and 40 ng/mL RANKL to generate osteoclasts. Osteoclast formation was completed by 6-8 days after RANKL treatment.

For tartrate resistant acid phosphatase (TRAP) staining, osteoclasts were rinsed in PBS and fixed with 4% paraformaldehyde for 20 minutes. The cells were then stained using Naphthol AS-MX phosphate and Fast Violet LB salt according to the protocol provided in BD Biosciences Technical Bulletin #445. Multinucleated (≥3 nuclei), TRAP-positive osteoclasts were counted in triplicate wells. Cells were imaged and photographed with light microscopy and the measurements were analyzed using NIH ImageJ (version 1.53).

The resorption activity of osteoclasts was examined using Osteo Assay Surface Plates (Corning) and cells were cultured as described above. Osteoclasts were removed by 5 minutes treatment with 10% bleach and washed with dH_2_O. The plates were then allowed to air dry completely at room temperature for 3–5 hours. The demineralized area was photographed by light microscopy and then analyzed using NIH ImageJ(^32^).

### MACROPHAGE ACTIVATION BY IFN-γ+LPS

BM derived macrophages (BMMs) were treated with IFN-γ at 100 units/ml overnight, followed by stimulation with 1 μg/mL *Escherichia coli*–derived LPS for 4 hours and 8 hours. Later, resting and IFN-γ+LPS–activated macrophages were subjected to RT-qPCR analysis.

### IMMUNOBLOTTING

Immunoblotting was performed as described previously(^16,25^). Briefly, cell protein lysates were harvested using a RIPA buffer (ThermoFisher Scientific) supplemented with Halt Protease & Phosphatase Inhibitor Cocktail (ThermoFisher Scientific). Protein concentration was determined using the Bradford assay. Equal amounts of protein (30-50 μg) were loaded on NuPAGE Bis-Tris 4%–12% gradient polyacrylamide gels in the MOPS or MES buffer system (ThermoFisher Scientific) and subsequently electrotransferred from mini gels to a solid nitrocellulose membrane using the iBlot 2 system (ThermoFisher Scientific). Immediately nitrocellulose membranes were saturated for at least 1 hour at room temperature in blocking buffer (LICOR Odyssey). The membranes were then incubated with appropriate primary antibody (0.1–1 μg/ml) in blocking buffer with 0.1% Tween-20 for overnight at 4°C. On the next day, the blot was incubated with anti-rabbit or anti-mouse HRP**−**conjugated secondary antibodies (Abcam) for 1 hour at room temperature. Antibody binding was detected using the ECL system (GE Healthcare). Images were acquired on G:BOX gel imaging system (Syngene) and cropped in Adobe Photoshop.

### QUANTITATIVE REAL-TIME PCR

The assay was performed as described previously(^16,25^). Briefly, total RNA was extracted from cultured cells using Trizol (Invitrogen). Complementary DNA was generated from 1 μg of total RNA with Superscript II RT (Invitrogen) and random hexamer primers (Promega). PCR was carried out by QuantStudio-3 Real-Time PCR System (Applied Biosystems) according to the manufacturer’s protocol. Transcript levels were normalized to *Hprt* and fold-changes were calculated by the 2^(-delta^ ^delta^ ^C(t))^ method(^33^). Primers used are summarized in **Table 2**.

**TABLE 2:**
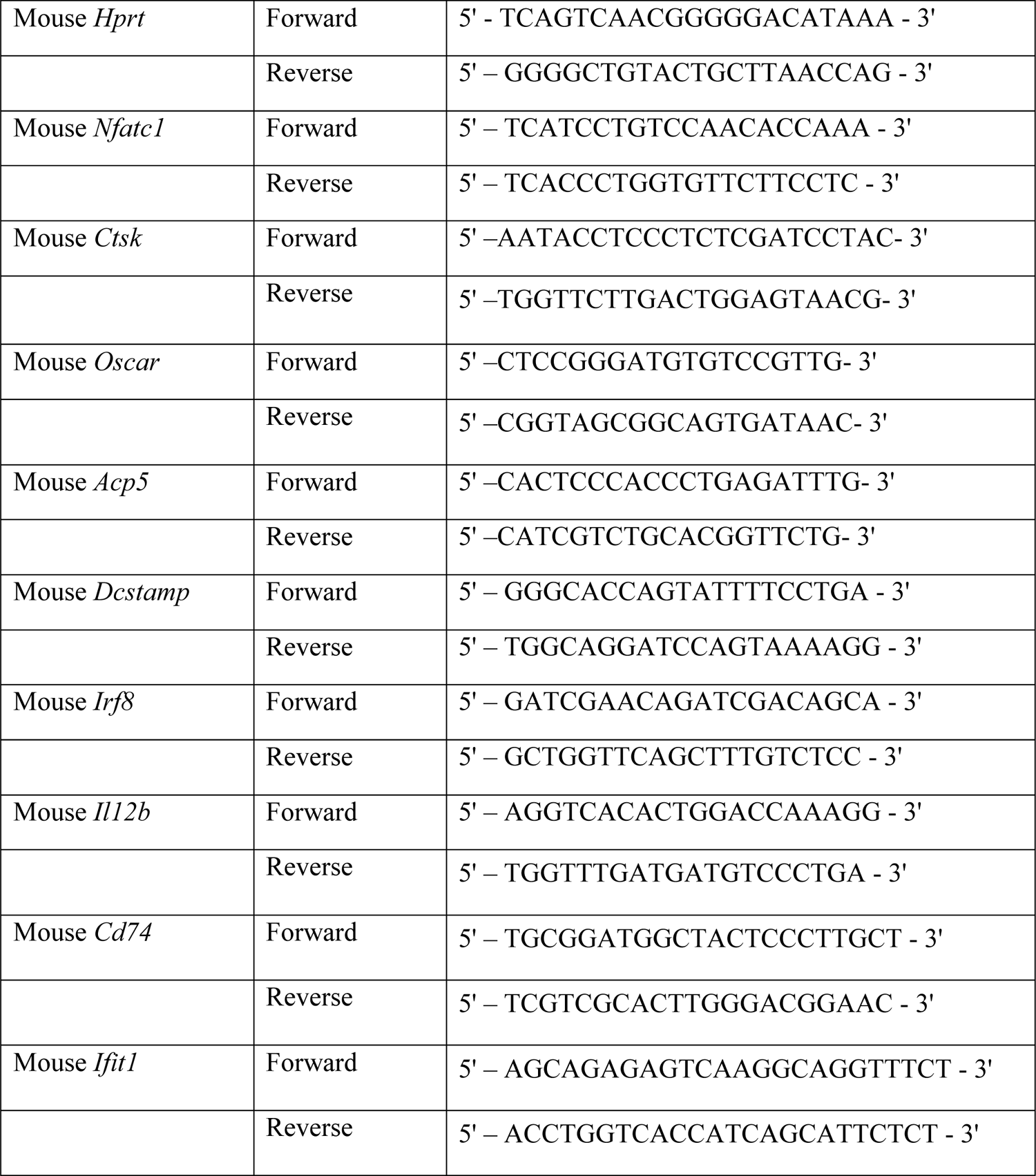
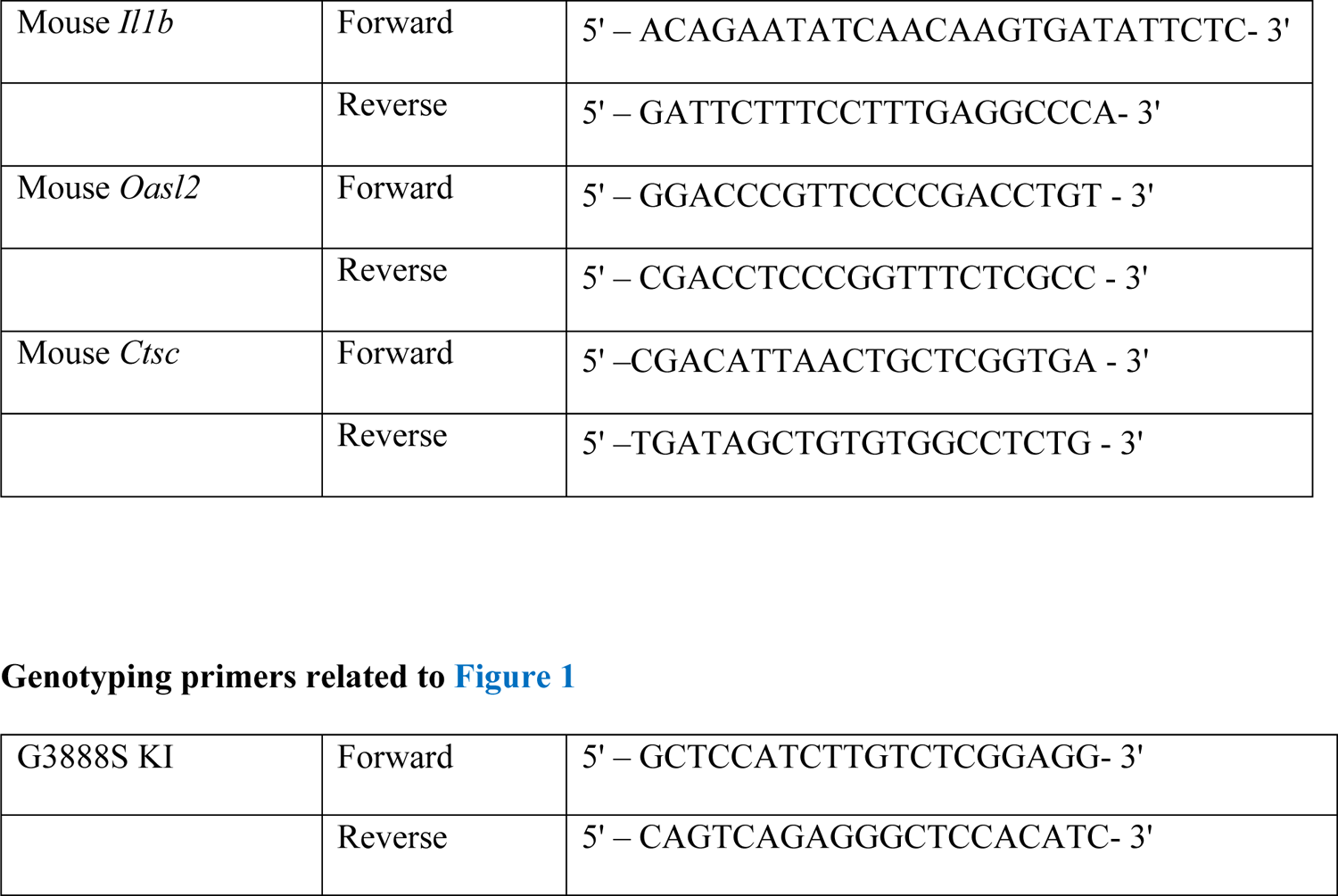
qPCR primers related to **Figures 3 and 5**

### ORAL LIGATURE MODEL

To investigate the *in vivo* effects of Irf8^G388S^ on osteoclast activity and tooth root resorption, a 5-0 silk suture (Roboz) was tied around the maxillary left second molar and left in place for 5 days(^34^). The contralateral maxillary molar tooth in each mouse was left unligated, which served as a baseline control for measurement of bone loss and root resorption. After five days, mice were euthanized and block biopsies from the maxillae were fixed in 10% formalin and then used to check bone loss and root resorption by micro–computed tomography (micro-CT) and histology.

### MICRO-CT ANALYSIS

Maxillae and femurs were scanned in a μCT 50 (Scanco Medical) at 70 kVp, 76 μA, 0.5 mm Al filter, with 900-1200 ms integration and 6 to 17 μm voxel dimension. DICOM files were created from raw data, exported, and calibrated to five known densities of hydroxyapatite (mg/cm^3^ HA). Reconstructed images were loaded and analyzed using AnalyzePro (version 14.0; AnalyzeDirect). For femur samples, calibrated images were reoriented and analyzed by standard cortical and trabecular bone algorithms(^16,35,36^). Maxillae were oriented to a uniform position using molar anatomical landmarks, and the region of interest (ROI) around the second maxillary molar was defined as the area between 240 µm distally and mesially of the most distal and mesial aspect of the second molar, respectively. Enamel was segmented at 1,600 mg/cm^3^ HA, while dentin/cementum and bone were segmented at 650 mg/cm^3^ HA, as previously described(^37^). We defined alveolar bone proper (ABP) as bone within 240 μm of the tooth root, as measured radially from tooth root surfaces to include buccal, lingual, radicular, and interproximal alveolar bone. Buccal bone was separated from lingual bone via subregional tracing using a line passing through the middle area of the furcation separating the buccal from lingual roots of the second maxillary molar. We calculated alveolar bone loss by subtracting the ligated areas (experimental) from non-ligated areas (control).

### HISTOLOGY

For histology, samples were fixed overnight in Bouin’s solution, hemisected, and demineralized in AFS (acetic acid, formaldehyde, sodium chloride) for 3-4 weeks and later embedded in paraffin, as previously described(^16,38^). Tartrate-resistant acid phosphatase (TRAP) staining was performed on decalcified and deparaffinized tissues to identify osteoclast/odontoclast-like cells according to manufacturer’s instructions (Wako Chemical, Japan)(^16^). For maxillae, two micrographs of molar roots were randomly selected from each animal for measurement of TRAP+ osteoclast and root resorption.

### QUANTIFICATION AND STATISTICAL ANALYSIS

Statistical tests were selected based on appropriate assumptions with respect to data distribution and variance characteristics. Data were analyzed by one-way ANOVA and post-hoc Tukey’s multiple comparison test using GraphPad Prism 6 (GraphPad Software, Inc.). Where appropriate (comparison of two groups only), two-tailed paired or unpaired Student’s *t* tests were performed. Throughout the paper, error bars indicate mean ± standard deviation (STD). *n* represents the number of biological replicates. For *in vitro* and *in vivo* experiments, number of biological replicates in each group is specified in the figure legends. Samples were excluded from analysis only in case of a clear technical problem that directly impacted endpoint measurements. Bone loss measurements and histological analysis were performed in a blinded fashion, whereas the conduct of experiments and assessment of other outcomes were not blinded to the study personnel. All *in vitro* experiments were performed three or more times (in triplicates) to ensure reproducibility of the observations.

## RESULTS

### Generation of *Irf8* KI Mice

To establish the molecular basis of increased osteoclast regulation and MICRR, we generated *Irf8* KI mice using CRISPR/Cas9 modeling the human *IRF8^G388S^* mutation (**Figure 1A**). The knock-in allele was validated by Sanger sequencing (**Figure 1B**) and PCR-based genotyping (**Figure 1C**). The *Irf8 KI* Het and Homo mice showed no evidence of gross developmental or morphological abnormalities compared to WT mice, in contrast to the *Irf8* global knockout (gKO) mice that are known to exhibit splenomegaly (**Figure 1D and E**). *Irf8 KI* Het and Homo mice were healthy, viable, and fertile, and exhibited no deviation from expected Mendelian ratios.

**Figure 1:**
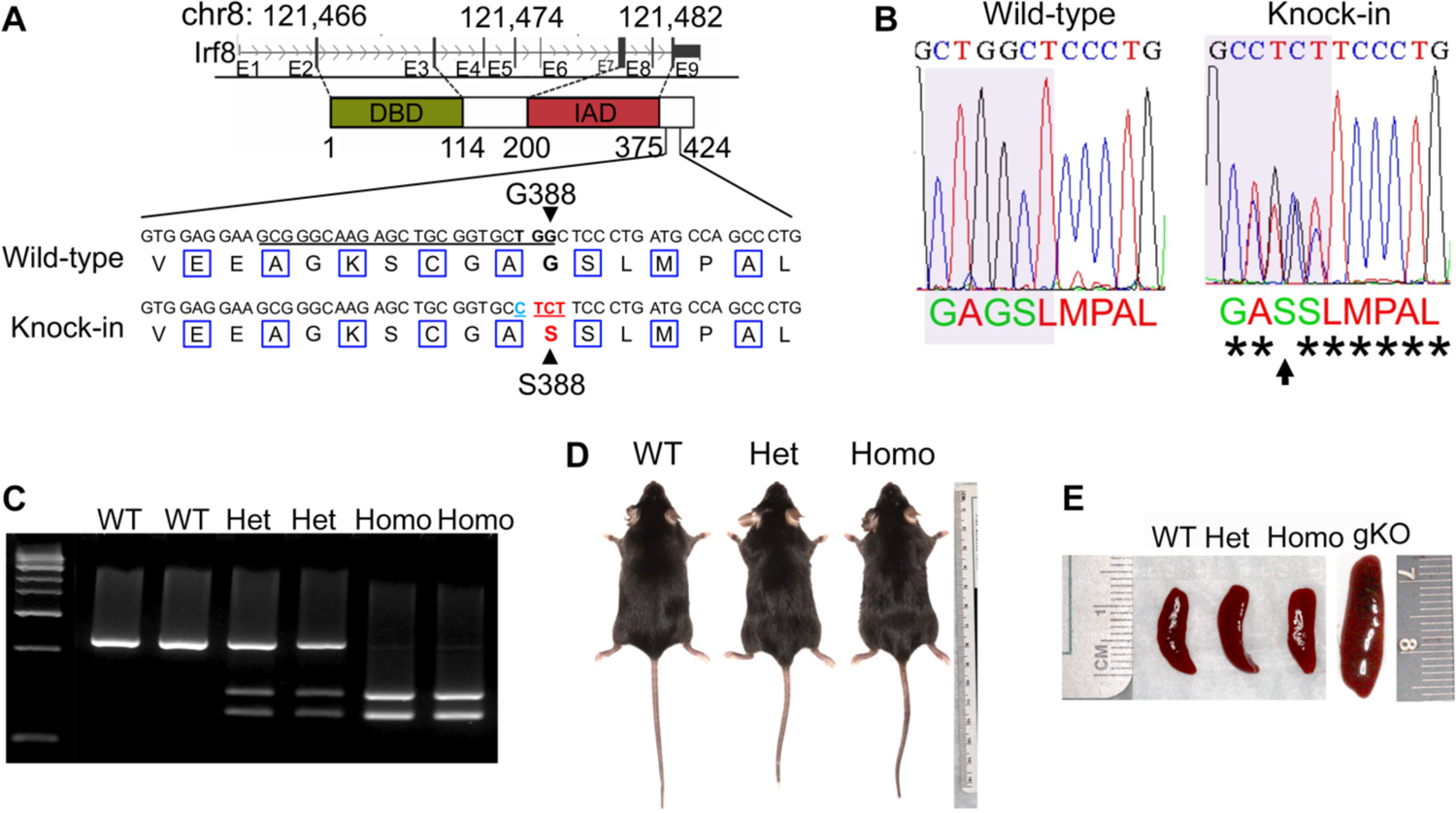
Generation of *Irf8* KI mice. (**A**) Schematic illustration of the G388S mutation in the mouse *Irf8* gene. Displayed at the top is the structure of the mouse *Irf8* gene, with introns represented by lines and the direction of transcription indicated by arrows. Exons are depicted as solid boxes, with coding regions as wide boxes while untranslated regions (UTRs) as narrow boxes. Directly beneath the gene structure is a diagram illustrating important domains of the IRF8 protein. The WT and mutant DNA sequences are listed at the bottom, with the CRISPR sgRNA binding region underlined and the PAM shown in bold. The altered amino acid and its codon are shown in red, while a silent mutation is shown in blue. The silent mutation does not result in amino acid change, but it can facilitate the using of PCR to differentiate the WT and KI alleles. (**B**) Sanger sequencing validated G388S knock-in allele in F2 mice. Left panel shows chromatograms of WT sequence. Right panel shows mutant reads. Gray shaded region indicates amino acid(s) at position 388 for each mouse. Arrow indicates desired knock-in mutation (G388S). (**C**) PCR of genomic DNA with primers flanking exons 2 and 3 shows that WT mice have one WT band (365-bp PCR product), heterozygous mice carry a WT band (365 bp) and two mutant bands (205 and 160 bp), and homozygous-null mice carry two mutant bands (205 and 160 bp). (**D**) Photographic images of representative WT, *Irf8* HET, and Homo mice aged 9-weeks-old. (**D**) Photographic images for spleen from *Irf8 KI* WT, HET, Homo, and gKO mice.

### *Irf8 KI* Mice Exhibit Unaltered Hematopoiesis

IRF8 is known to enforce monocyte development by opposing neutrophil lineage(^39,40^), and additionally promoting the differentiation of common lymphoid progenitors (CLPs) to B cells(^41,42^), driving effector differentiation of CD8 T cells(^43^), and regulating the development of type 1 conventional DCs (cDC1s) and plasmacytoid DCs (pDCs)(^18,40,44–46^). Consequently, humans and mice harboring mutations within the IRF8 DNA-binding domain and IAD domain exhibit reduced monocytes, B-cells, T-cells, cDCs, and pDCs, and are immunodeficient(^18,40,42,44–46^). The development of myeloid and lymphoid lineage cells in *Irf8 KI* Het and Homo mice was examined by flow cytometric analysis by comparing BM, blood, and spleen cells for expression of cell lineage markers. In contrast to the above-mentioned findings, we noted that the development of monocytes, neutrophils, B-cells, and T-cells were normal in *Irf8* KI Het and Homo mice when compared to WT mice (**Figure 2A-C)**. These results are consistent with MICRR patients harboring the *IRF8^G388S^* mutation(^16^) who showed no deficiencies in immune cells, suggesting that the G388S variant does not negatively impact hematopoietic cell development.

**Figure 2:**
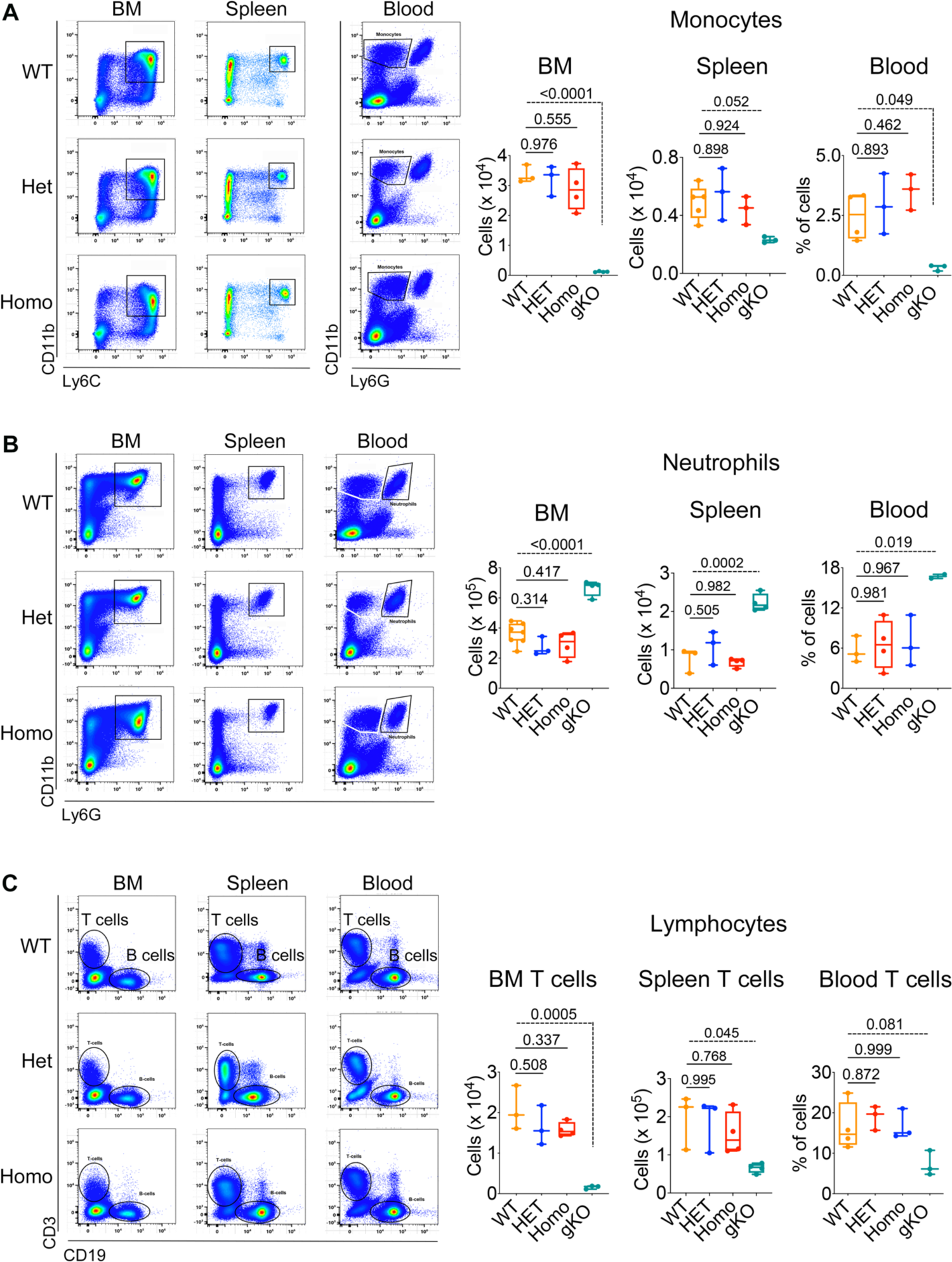
*Irf8^G388S^* has no impact on the development of myeloid and lymphoid lineage cells. Flow cytometric analysis of (**A**) monocytes, (**B**) neutrophils, and (**C**) lymphocytes in BM, blood, and spleen. Pseudocolor plots show cell population and bar graphs show absolute counts. Data are representative of at least 3-4 independent experiments. The data are presented as the mean ± STD (n = 3-4 mice per genotype). One-way ANOVA and post-hoc Tukey’s test was used for comparisons among groups. Cells were gated as previously described(^25^). The detailed results of *Irf8* gKO mice has been reported elsewhere(^25^).

### *Irf8 KI* Mice Exhibit Normal Macrophage Function

IRF8, along with IRF1 and PU.1, is known to regulate the expression of genes important for macrophage function(^47–49^). Hence, IRF8 deficiency renders macrophages hypofunctional in their response to IFN-γ(^50^), production of inflammatory cytokines(^47^), and defense against intracellular infections(^21,48^). In order to determine the effects of *Irf8^G388S^* mutation on macrophage function, we performed RT-qPCR analysis on resting and IFN-γ+LPS–activated macrophages. IFN-γ caused upregulation of several macrophage-related genes, which were further amplified upon LPS stimulation (**Figure 3**), however, there were no significant differences between *Irf8* KI Het and Homo mice versus WT mice. These findings suggest that the *Irf8^G388S^* variant has minimal impact on macrophage gene signatures that are important for antimicrobial defenses and inflammatory cytokine production.

**Figure 3:**
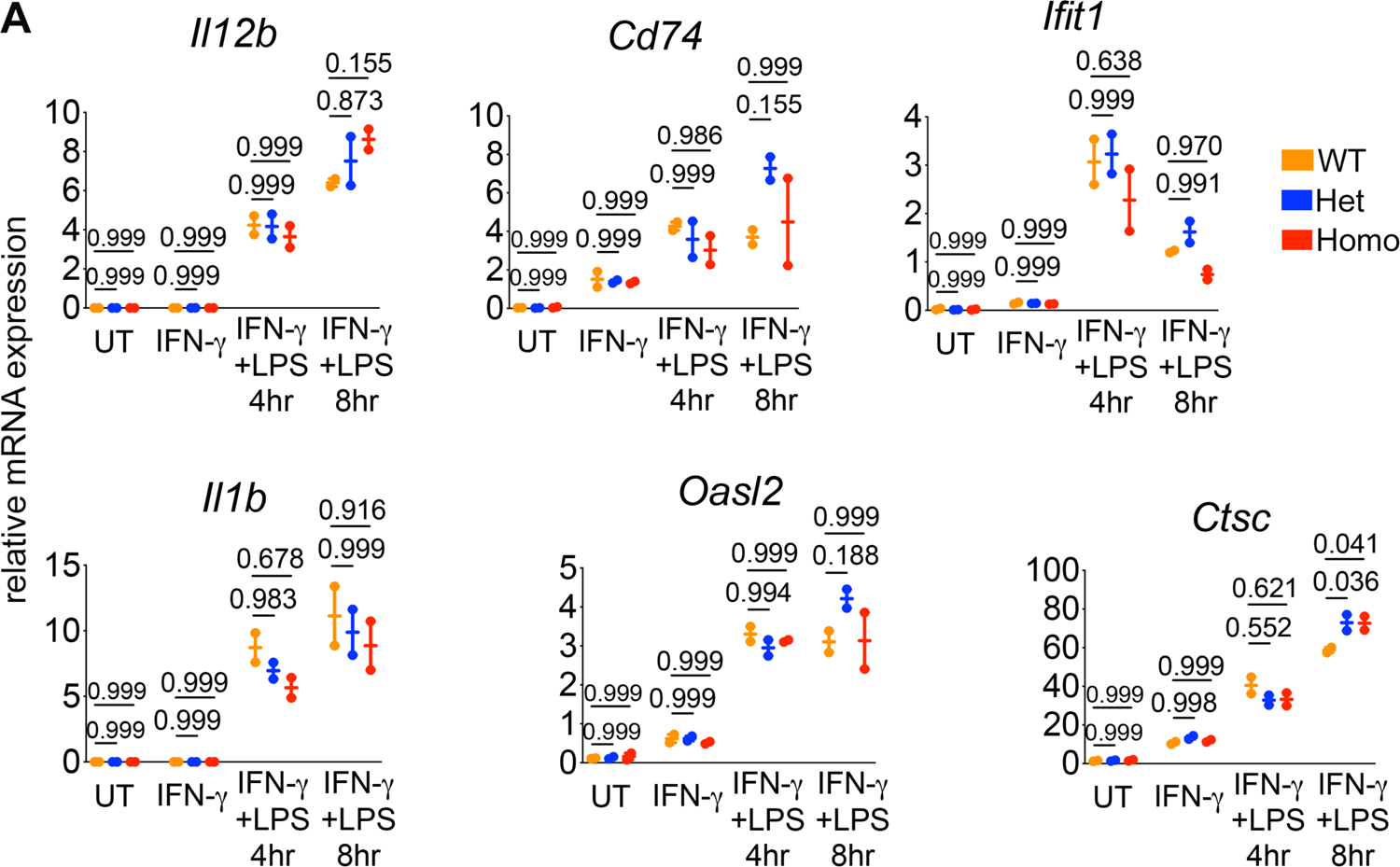
*Irf8^G388S^* has minimal impact on macrophage gene signatures important for antimicrobial defenses and inflammatory cytokine production. **(A)** RT-qPCR analysis of resting and IFN-γ+LPS–activated macrophages. Data are representative of at least three independent experiments. The data are presented as the mean ± STD (n = 3 mice per genotype). One-way ANOVA and post-hoc Tukey’s test was used for comparisons among groups.

### *Irf8 KI* Mice Exhibit Increased Osteoclastogenesis

IRF8 deficiency is known to promote increased osteoclastogenesis(^16,23,25^). Hence, to determine the effects of *Irf8^G388S^* mutation on osteoclast regulation, we examined *Irf8 KI* Het and Homo mice for abnormal bone and OC phenotypes. We noted that *Irf8 KI* Het and Homo mice respectively displayed reduced bone mass accompanied by dramatic decreases in trabecular bone mineral density (29% and 28%), cortical thickness (16% and 18%), and cortical area fraction (12% and 13%) when compared to WT mice (**Figure 4**). To measure osteoclast formation and its activity *in vitro*, we cultured BMMs from WT and *Irf8 KI* mice with M-CSF and RANKL for 6-8 days. Approximately 1.5 to 2-fold increased number of osteoclasts formed in *Irf8 KI* Het and Homo cultures with cell size ∼1.5-fold larger than WT cells, which was appended by a 2 to 3-fold increase in resorption activity when compared to WT cells (**Figure 5A**). Correspondingly, the mRNA and protein expression of NFATc1 and its downstream osteoclast-related genes such as *Ctsk*, *Acp5*, and *Dcstamp* were significantly upregulated in both *Irf8 KI* Het and Homo osteoclasts when compared to WT osteoclasts (**Figure 5B and 5C**). Taken together, these results establish that the loss of IRF8 regulatory function in *Irf8 KI* mice due to G388S mutation promotes increased osteoclastogenesis.

**Figure 4:**
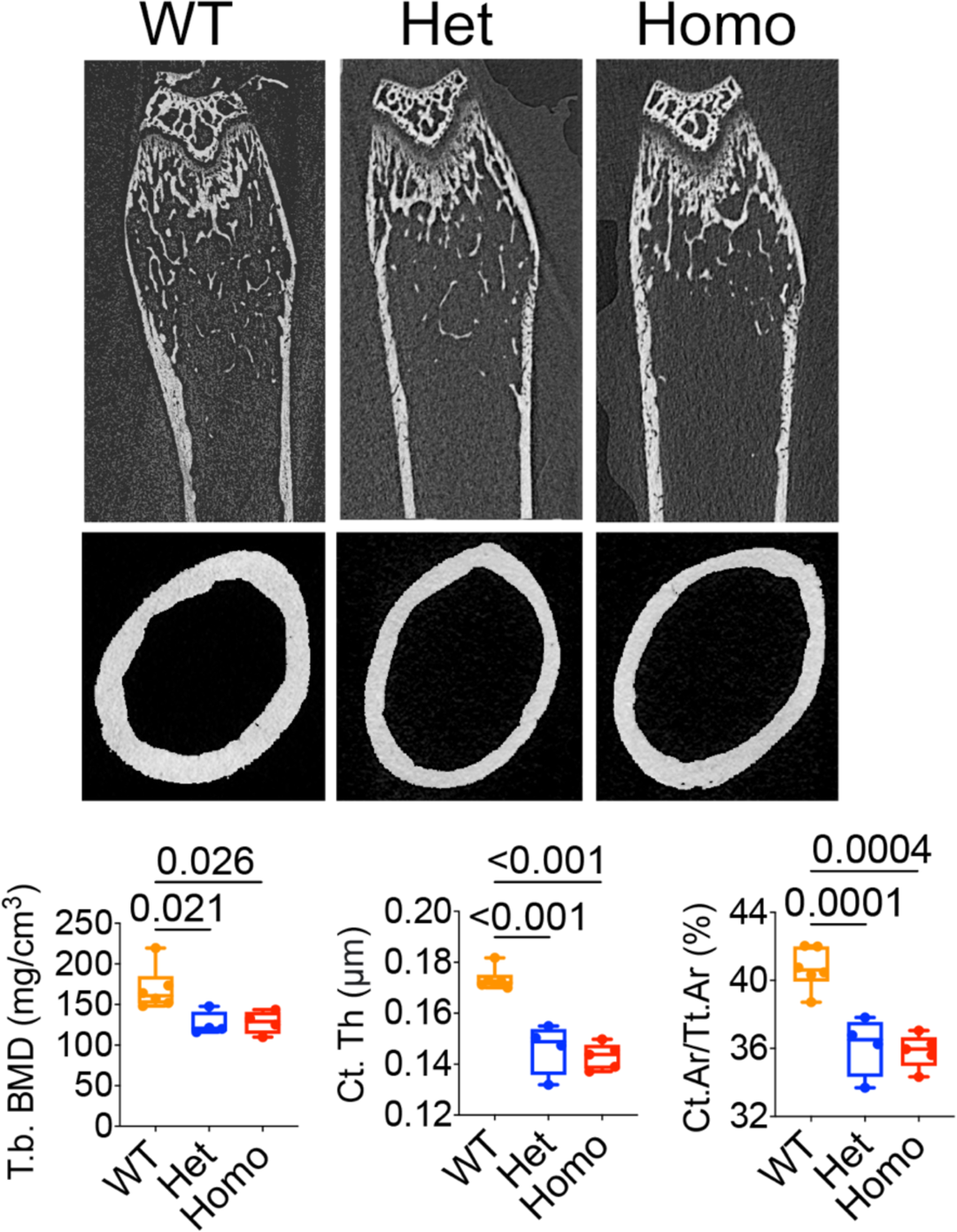
*Irf8^G388S^* promotes increased osteoclast activity in femur. **(A)** Micro-CT analysis of femurs (9-week-old mice). Top, longitudinal view; middle, axial view of the cortical bone in midshaft; bottom, axial view of the trabecular bone in metaphysis. Scale bar, 0.5 mm. Bar graphs show bone morphometric analysis of femurs. Tb. BMD, trabecular bone mineral density; Ct. Th, cortical thickness; Ct.Ar/Tt.Ar, cortical area fraction. n = 4-6 mice per genotype. The data are presented as the mean ± STD. One-way ANOVA and post-hoc Tukey’s test was used for comparisons among groups.

**Figure 5:**
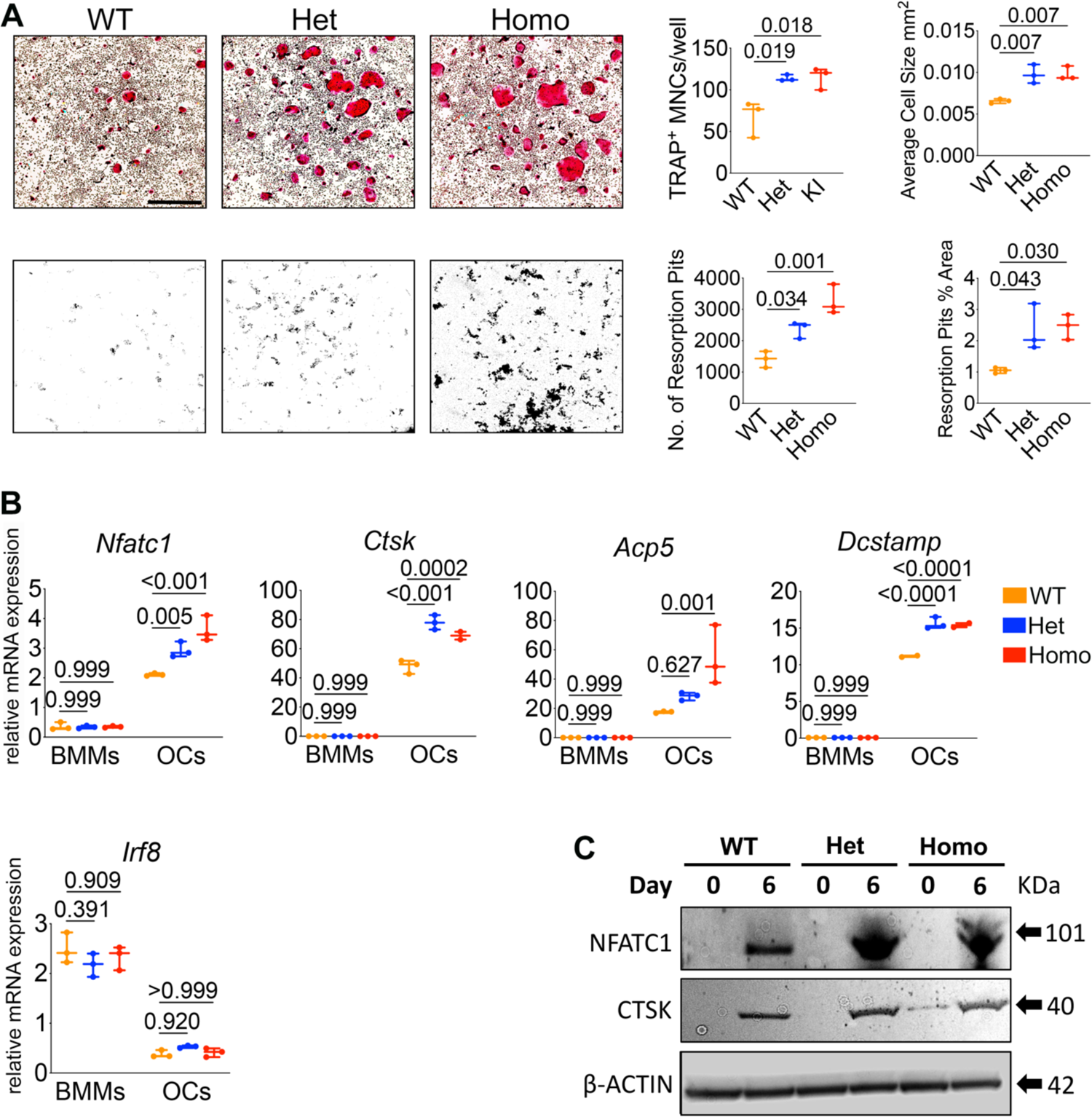
*Irf8^G388S^* promotes increased in vitro osteoclastogenesis. **(A)** Top panel, TRAP-stained cells show OC formation. Bottom panel shows pit formation ability of OCs. Scale bar, 1000 μm. Bar graphs show quantified number of TRAP^+^ cells, average cell size of TRAP^+^ cells, resorption pits, and percentage area of resorption in each group. **(B)** RT-qPCR analysis of OC-specific genes in BMMs and OCs. **(C)** Immunoblot analysis of OC-specific proteins. Day 0 = BMMs and Day 6 = OCs. (D-F) Data are representative of at least three independent experiments performed in triplicates. The data are presented as the mean ± STD. One-way ANOVA and post-hoc Tukey’s test was used for comparisons among groups.

### *Irf8^G388S^* Mutation Fails to Inhibit NFATc1 Transcriptional Activity

The IRF8 IAD domain (IAD1: N/200-377aa) is known to physically interact with the TAD-A domain of NFATc1 (N/1-205aa) to inhibit NFATc1 nuclear translocation and activation of downstream target genes(^28^). The identified G388S mutation is 11 amino acids downstream of the IAD domain, which could impact IRF8 interaction with NFATc1. To gain insights into the functional consequences of hIRF8^G388S^ mutation on osteoclastogenesis, we created expression vectors of human IRF8 wild type (hIRF8^WT^) and G388S mutant (hIRF8^G388S^) and examined their effects on the transcriptional activation and physical interaction with NFATc1, the results of which have been reported in detail elsewhere(^16^). Briefly, overexpression of *Nfatc1* gene activated *Ctsk* promoter, while simultaneous expression of hIRF8^WT^ reduced the activation of *Ctsk* promoter, indicating that hIRF8^WT^ inhibits the transcriptional activity of NFATc1. Conversely, hIRF8^G388S^ mutant failed to inhibit NFATc1-dependent transcriptional activation of *Ctsk* promoter, suggesting that the mutation disrupts functional binding of IRF8 with NFATc1. Coimmunoprecipitation assay confirmed that hIRF8^G388S^ failed to physically interact with NFATc1 when compared to hIRF8^WT^(^16^). Protein stability assay further showed that hIRF8^G388S^ protein was stable and that the loss of IRF8 function is more likely because of its inability to interact with other transcription factors such as the NFATc1(^16^).

### Oral Ligature Inserted *Irf8 KI* Mice Exhibit Increased Alveolar Bone Loss and Root Resorption

To investigate the effects of G388S mutation on *in vivo* osteoclast activity in the dentoalveolar region, we utilized the oral ligature model that is known to promote oral bacteria-mediated inflammatory bone loss(^34^). At a steady state, we noted no root resorption activity in *Irf8 KI* Het and Homo mice when compared to WT mice. Following ligature placement around maxillary 2^nd^ molar for 5 days, histological analyses of maxillae showed increased number of osteoclasts lining the alveolar bone (surrounding molar teeth) in *Irf8 KI* Het and Homo mice when compared to WT mice (**Figure 6A**). Increased osteoclast numbers were consequently associated with increased alveolar bone resorption in *Irf8 KI* Het and Homo mice when compared to WT mice (**Figure 6A**). Micro-CT analysis corroborated increased alveolar bone loss in *Irf8 KI* Het and Homo mice when compared to WT mice (**Figure 6B**). Similarly, we noted we noted increased number of osteoclasts lining the root surface of 2^nd^ molar tooth leading to increased root resorption in *Irf8 KI* Het and Homo mice when compared to WT mice (**Figures 6A**). The increased osteoclast activity in the dentoalveolar region is directly correlated with increased NFATc1 expression in *Irf8* deficient mice(^16^). Taken together, these results suggest that the increased osteoclast activity noted in *Irf8 KI* Het and Homo mice is a consequence of loss of *Irf8* function, which supports an important parallel to our patients with heterozygous *IRF8* G388S mutation and increased frequency of osteoclast/odontoclast activity in the jaw bones.

**Figure 6:**
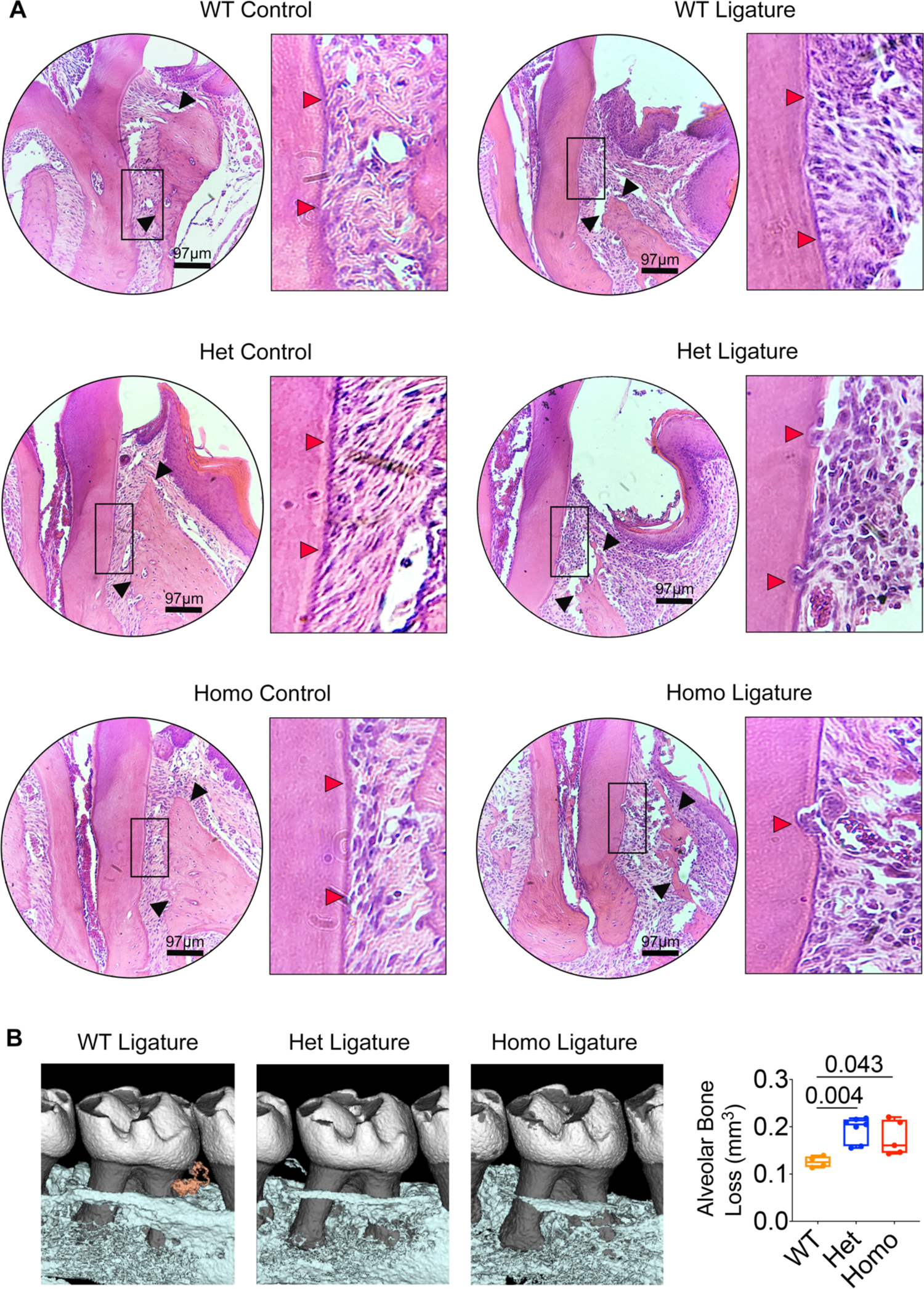
*Irf8^G388S^* promotes increased alveolar bone loss and tooth root resorption. **(A)** Histological analyses of maxillae (9-week-old mice) by H&E staining demonstrate increased alveolar bone loss (black arrows) and tooth root resorption (red arrows) in *Irf8 KI* Het and Homo mice compared to WT mice. **(B)** Micro-CT analyses of maxillae. 2D cut planes in sagittal orientation shows alveolar bone loss around ligated maxillary 2^nd^ molar. Scale bar, 0.5 mm. Bar graphs show quantified results for alveolar bone loss around maxillary 2^nd^ molar in each group. n = 5-6 mice per genotype. The data are presented as the mean ± STD. One-way ANOVA and post-hoc Tukey’s test was used for comparisons among groups.

## DISCUSSION

The *Irf8^G388S^ KI* mice generated in this study provide a unique translational model to study (1) the etiopathology of MICRR, (2) the critical role of IRF8 in osteoclast regulation, (3) the IRF8 domain important for osteoclast function, and (4) develop targeted therapies for MICRR and other skeletal disorders mediated by increased osteoclast activity.

To date, several *IRF8* mutations have been identified in humans that cause a range of immune cell phenotypes leading to increase susceptibility to infectious diseases(^51–54^). The homozygous K108E variant, located in the DNA-binding domain, causes severe immunodeficiency and a complete lack of monocytes, DCs, IL-12, IFN-γ and TNF-α production(^51,52^). On the other hand, the heterozygous T80A variant, located in the key DNA-binding helix, is associated with milder immunodeficiency and selective depletion of the CD1c+ compartment of CD11c+ circulating DCs(^51^). The compound heterozygous mutations R83C/R291Q cause a complex immunodeficiency syndrome characterized by DC and monocyte deficiency(^53^). The biallelic A201V/P224L mutation within the IAD domain leads to NK cell deficiency, decreased NK cell number, and CD56^dim^ subset(^54^). However, unlike the K108E mutation, the A201V/P224L variant can activate IRF1- and PU.1-dependent transcription normally, which results in a subtle DC deficiency and normal production of IFN-γ and TNF-α. Together, these findings highlight the varying impact of different IRF8 mutations on immune cells. However, there are no reports of detailed dental or skeletal phenotypes in patients with these mutations, except for a history of oral candidiasis and normal osteoclast activity in a 10-week-old infant carrying the K108E variant(^51^).

In contrast to these previously reported mutations, we show that the *Irf8^G388S^* mutation mainly affects osteoclastogenesis, sparing immune cell development and function. The mechanisms underlying the differential impact of *IRF8* mutations on immune cells vs. osteoclasts remain unclear. IRF8 by itself possesses weak transcriptional activity. However, it can function both as a transcriptional activator and repressor by forming different DNA-binding heterocomplexes with multiple partners, including IRF, ETS and NFAT family members(^26–29^). It is plausible that variants in the DNA-binding domain (DBD) vs. IRF-association domain (IAD) may differentially affect IRF8 heterodimerization with other partners, leading to differing phenotypes. The IRF8 IAD domain is known to physically interact with the TAD-A domain of NFATc1 to inhibit NFATc1 nuclear translocation and activation of downstream target genes(^28^). We show that the mutant IRF8 G388S isoform fails to physically interact with NFATc1 and negatively inhibit NFATc1-dependent transcriptional activation, thus leading to increased osteoclastogenesis(^16^). The interaction of mutant IRF8 G388S isoform with other transcription partners, such as ETS and IRF family members needs to be investigated.

Currently, the only other mutation associated with MICRR is a missense mutation c.5630 C > T in the filamin A (*FLNA*) gene located on the chromosome X, suggesting the possibility of sex-linked recessive inheritance(^55^). The 19-year-old male inherited the mutated X chromosome from his mother and reported non-contributory medical and dental history with laboratory results within normal ranges(^55^). The *FLNA* gene encodes for actin-binding protein or filamin that crosslinks actin filaments into orthogonal networks in the cortical cytoplasm and plays a role in anchoring membrane proteins to the actin cytoskeleton(^56,57^). FLNA is required for monocyte migration during osteoclastogenesis via its role in regulating Rho GTPase-mediated actin remodeling(^58,59^). FLNA also interacts with low-density lipoprotein receptor-related proteins 6 (LRP6) to inhibit β-catenin expression and enhances NFATc1-dependent osteoclastogenic gene expression to inhibit osteogenesis and promote osteoclastogenesis(^60^). Based on these findings and previously reported “gain-of-function” *FLNA* mutations that are known to cause congenital malformations affecting craniofacial and skeletal structures, the authors speculate that the *FLNA* c.5630 C > T mutation possibly leads to augmented odontoclastic/osteoclastic activity(^55^). However, no functional assays were performed to validate this hypothesis. Empirical observations suggest that MICRR may not be a monogenic disease, and other regulators upstream or downstream of the osteoclast signaling pathway could potentially contribute to root resorption.

In our study, the *Irf8 KI* Het and Homo mice exhibited increased alveolar bone loss and root resorption following oral ligature insertion. At a steady state, we noted no tooth root resorption activity in these animals when compared to WT mice. These findings suggest that IRF8 dysfunction, increased osteoclast activity, and oral microbiota collectively contribute to the development of root resorption in mice. Similarly, in affected individuals, increased osteoclast activity alone wouldn’t be sufficient to promote MICRR. Patient specific factors such as developmental tooth defects, oral microbiome, trauma, bruxism, and environmental exposure are needed to collectively increase susceptibility to MICRR. MICRR is not always noted in patients with overactive osteoclast-related disorders, further supporting that MICRR is a multifactorial disease and involves etiological factors specific to the oral cavity. This notion further helps explain why patients with *IRF8* G388S mutation and *FLNA* c.5630 C > T mutation mainly exhibited dental problems and no other obvious systemic bone disorders.

In summary, this translational study delineates the IRF8 domain important for osteoclast function and provides novel insights into the etiopathology of MICRR and *IRF8* mutation associated with MICRR. The IRF8 IAD domain appears to be more important for osteoclast function vs immune cell function. Dysregulation of osteoclast differentiation/activity appears to be the ‘common denominator’ or ‘central triggering issue’ for MICRR, further complemented by animal/patient specific factors. Agents that modulate osteoclast activity, such as bisphosphonates and RANKL antibodies, may prove beneficial in preventing and treating MICRR, although the risks of medication-related osteonecrosis of the jaw (MRONJ) associated with these anti-resorptive medications must be carefully considered and investigated further.

## Acknowledgements

We thank Dr. Anthony Neely from Univ. of Detroit-Mercy for data related to research subjects. We thank Dr. Xiaoxuan Fan from Univ. of Maryland for assistance with flow cytometry. We thank Dr. Martha Somerman from National Institutes of Health for reviewing and critically editing the manuscript.

## AUTHOR CONTRIBUTIONS

V.T.M., A.D., and S.Y., conceived the project and designed the experiments; C.L., helped with generating KI mice using CRISPR/Cas9; A.D., S.Y., X.W., J.G., H.T., B.K., and V.T.M., performed the experiments; S.W., and D.A., inserted oral ligatures in mice; S.Y., and H.T., performed flow cytometry analysis; C.Z., and B.L.F., performed micro-CT analysis; K.O., B.L.F., and M.C., advised on experiments and provided intellectual input; V.T.M., wrote the paper, supervised the project; and all authors contributed to reviewing and editing.

## REFERENCES

1. Harokopakis-Hajishengallis E. Physiologic root resorption in primary teeth: molecular and histological events. Journal of oral science. Mar 2007;49(1):1–12.

2. Darcey J, Qualtrough A. Resorption: part 2. Diagnosis and management. British dental journal. May 2013;214(10):493–509.

3. Darcey J, Qualtrough A. Resorption: part 1. Pathology, classification and aetiology. British dental journal. May 2013;214(9):439–51.

4. Fuss Z, Tsesis I, Lin S. Root resorption--diagnosis, classification and treatment choices based on stimulation factors. Dental traumatology: official publication of International Association for Dental Traumatology. Aug 2003;19(4):175–82.

5. Ne RF, Witherspoon DE, Gutmann JL. Tooth resorption. Quintessence international. Jan 1999;30(1):9–25.

6. Liang H, Burkes EJ, Frederiksen NL. Multiple idiopathic cervical root resorption: systematic review and report of four cases. Dento maxillo facial radiology. May 2003;32(3):150–5.

7. Beckett HA, Gilmour AG. Multiple idiopathic cervical root resorption in a male. British dental journal. Jul 10 1993;175(1):33–4.

8. Wu J, Lin LY, Yang J, Chen XF, Ge JY, Wu JR, et al. Multiple idiopathic cervical root resorption: a case report. International endodontic journal. 2015:n/a-n/a.

9. Yu VS, Messer HH, Tan KB. Multiple idiopathic cervical resorption: case report and discussion of management options. International endodontic journal. Jan 2011;44(1):77–85.

10. Iwamatsu-Kobayashi Y, Satoh-Kuriwada S, Yamamoto T, Hirata M, Toyoda J, Endo H, et al. A case of multiple idiopathic external root resorption: a 6-year follow-up study. Oral surgery, oral medicine, oral pathology, oral radiology, and endodontics. Dec 2005;100(6):772–9.

11. Jiang YH, Lin Y, Ge J, Zheng JW, Zhang L, Zhang CY. Multiple idiopathic cervical root resorptions: report of one case with 8 teeth involved successively. International journal of clinical and experimental medicine. 2014;7(4):1155–9.

12. Macdonald-Jankowski D. Multiple idiopathic cervical root resorption most frequently seen in younger females. Evidence-based dentistry. 2005;6(1):20.

13. Neely AL, Gordon SC. A familial pattern of multiple idiopathic cervical root resorption in a father and son: a 22-year follow-up. Journal of periodontology. Feb 2007;78(2):367–71.

14. Neely AL, Thumbigere-Math V, Somerman MJ, Foster BL. A Familial Pattern of Multiple Idiopathic Cervical Root Resorption With a 30-Year Follow-Up. Journal of periodontology. Apr 2016;87(4):426–33. Epub 2015/11/13.

15. Chu EY, Deeb JG, Foster BL, Hajishengallis E, Somerman MJ, Thumbigere-Math V. Multiple Idiopathic Cervical Root Resorption: A Challenge for a Transdisciplinary Medical-Dental Team. Front Dent Med. Mar 2021;2. Epub 2021/08/10.

16. Thumbigere-Math V, Foster BL, Bachu M, Yoshii H, Brooks SR, Coulter A, et al. Inactivating Mutation in IRF8 Promotes Osteoclast Transcriptional Programs and Increases Susceptibility to Tooth Root Resorption. J Bone Miner Res. Jun 2019;34(6):1155–68. Epub 2019/03/07.

17. Tamura T, Ozato K. ICSBP/IRF-8: its regulatory roles in the development of myeloid cells. Journal of interferon & cytokine research: the official journal of the International Society for Interferon and Cytokine Research. Jan 2002;22(1):145–52.

18. Aliberti J, Schulz O, Pennington DJ, Tsujimura H, Reis e Sousa C, Ozato K, et al. Essential role for ICSBP in the in vivo development of murine CD8alpha + dendritic cells. Blood. Jan 1 2003;101(1):305–10.

19. Tailor P, Tamura T, Morse HC 3rd, Ozato K. The BXH2 mutation in IRF8 differentially impairs dendritic cell subset development in the mouse. Blood. Feb 15 2008;111(4):1942-5.

20. Tamura T, Nagamura-Inoue T, Shmeltzer Z, Kuwata T, Ozato K. ICSBP directs bipotential myeloid progenitor cells to differentiate into mature macrophages. Immunity. Aug 2000;13(2):155–65.

21. Marquis JF, LaCourse R, Ryan L, North RJ, Gros P. Disseminated and rapidly fatal tuberculosis in mice bearing a defective allele at IFN regulatory factor 8. Journal of immunology. Mar 1 2009;182(5):3008–15.

22. Turcotte K, Gauthier S, Malo D, Tam M, Stevenson MM, Gros P. Icsbp1/IRF-8 is required for innate and adaptive immune responses against intracellular pathogens. Journal of immunology. Aug 15 2007;179(4):2467–76.

23. Zhao B, Takami M, Yamada A, Wang X, Koga T, Hu X, et al. Interferon regulatory factor-8 regulates bone metabolism by suppressing osteoclastogenesis. Nature medicine. Sep 2009;15(9):1066–71.

24. Fang C, Qiao Y, Mun SH, Lee MJ, Murata K, Bae S, et al. Cutting Edge: EZH2 Promotes Osteoclastogenesis by Epigenetic Silencing of the Negative Regulator IRF8. Journal of immunology. Jun 1 2016;196(11):4452–6. Epub 2016/05/18.

25. Das A, Wang X, Kang J, Coulter A, Shetty AC, Bachu M, et al. Monocyte Subsets With High Osteoclastogenic Potential and Their Epigenetic Regulation Orchestrated by IRF8. J Bone Miner Res. Jan 2021;36(1):199–214. Epub 2020/08/18.

26. Sharf R, Azriel A, Lejbkowicz F, Winograd SS, Ehrlich R, Levi BZ. Functional domain analysis of interferon consensus sequence binding protein (ICSBP) and its association with interferon regulatory factors. The Journal of biological chemistry. Jun 2 1995;270(22):13063–9.

27. Bovolenta C, Driggers PH, Marks MS, Medin JA, Politis AD, Vogel SN, et al. Molecular interactions between interferon consensus sequence binding protein and members of the interferon regulatory factor family. Proceedings of the National Academy of Sciences of the United States of America. May 24 1994;91(11):5046–50.

28. Jiang DS, Wei X, Zhang XF, Liu Y, Zhang Y, Chen K, et al. IRF8 suppresses pathological cardiac remodelling by inhibiting calcineurin signalling. Nature communications. 2014;5:3303. Epub 2014/02/15.

29. Kanno Y, Levi BZ, Tamura T, Ozato K. Immune cell-specific amplification of interferon signaling by the IRF-4/8-PU.1 complex. Journal of interferon & cytokine research: the official journal of the International Society for Interferon and Cytokine Research. Dec 2005;25(12):770-9.

30. Wang H, Yang H, Shivalila CS, Dawlaty MM, Cheng AW, Zhang F, et al. One-step generation of mice carrying mutations in multiple genes by CRISPR/Cas-mediated genome engineering. Cell. May 9 2013;153(4):910–8. Epub 2013/05/07.

31. Cossarizza A, Chang HD, Radbruch A, Acs A, Adam D, Adam-Klages S, et al. Guidelines for the use of flow cytometry and cell sorting in immunological studies (second edition). Eur J Immunol. Oct 2019;49(10):1457–973. Epub 2019/10/22.

32. Schneider CA, Rasband WS, Eliceiri KW. NIH Image to ImageJ: 25 years of image analysis. Nat Methods. Jul 2012;9(7):671–5. Epub 2012/08/30.

33. Livak KJ, Schmittgen TD. Analysis of relative gene expression data using real-time quantitative PCR and the 2(-Delta Delta C(T)) Method. Methods. Dec 2001;25(4):402–8. Epub 2002/02/16.

34. Abe T, Hajishengallis G. Optimization of the ligature-induced periodontitis model in mice. J Immunol Methods. Aug 30 2013;394(1-2):49–54. Epub 2013/05/16.

35. Bouxsein ML, Boyd SK, Christiansen BA, Guldberg RE, Jepsen KJ, Muller R. Guidelines for assessment of bone microstructure in rodents using micro-computed tomography. J Bone Miner Res. Jul 2010;25(7):1468–86. Epub 2010/06/10.

36. Thumbigere-Math V, Alqadi A, Chalmers NI, Chavez MB, Chu EY, Collins MT, et al. Hypercementosis Associated with ENPP1 Mutations and GACI. Journal of dental research. Apr 2018;97(4):432–41. Epub 2017/12/16.

37. Chavez MB, Chu EY, Kram V, de Castro LF, Somerman MJ, Foster BL. Guidelines for Micro-Computed Tomography Analysis of Rodent Dentoalveolar Tissues. JBMR Plus. Mar 2021;5(3):e10474. Epub 2021/03/30.

38. Foster BL. Methods for studying tooth root cementum by light microscopy. Int J Oral Sci. Sep 2012;4(3):119–28. Epub 2012/09/22.

39. Kurotaki D, Yamamoto M, Nishiyama A, Uno K, Ban T, Ichino M, et al. IRF8 inhibits C/EBPalpha activity to restrain mononuclear phagocyte progenitors from differentiating into neutrophils. Nature communications. Sep 19 2014;5:4978. Epub 2014/09/23.

40. Becker AM, Michael DG, Satpathy AT, Sciammas R, Singh H, Bhattacharya D. IRF-8 extinguishes neutrophil production and promotes dendritic cell lineage commitment in both myeloid and lymphoid mouse progenitors. Blood. Mar 1 2012;119(9):2003–12. Epub 2012/01/13.

41. Wang H, Lee CH, Qi C, Tailor P, Feng J, Abbasi S, et al. IRF8 regulates B-cell lineage specification, commitment, and differentiation. Blood. Nov 15 2008;112(10):4028–38.

42. Wang H, Jain S, Li P, Lin JX, Oh J, Qi C, et al. Transcription factors IRF8 and PU.1 are required for follicular B cell development and BCL6-driven germinal center responses. Proceedings of the National Academy of Sciences of the United States of America. Apr 18 2019. Epub 2019/04/20.

43. Miyagawa F, Zhang H, Terunuma A, Ozato K, Tagaya Y, Katz SI. Interferon regulatory factor 8 integrates T-cell receptor and cytokine-signaling pathways and drives effector differentiation of CD8 T cells. Proceedings of the National Academy of Sciences of the United States of America. Jul 24 2012;109(30):12123–8. Epub 2012/07/12.

44. Schiavoni G, Mattei F, Sestili P, Borghi P, Venditti M, Morse HC, 3rd, et al. ICSBP is essential for the development of mouse type I interferon-producing cells and for the generation and activation of CD8alpha(+) dendritic cells. J Exp Med. Dec 2 2002;196(11):1415-25. Epub 2002/12/04.

45. Tsujimura H, Tamura T, Ozato K. Cutting edge: IFN consensus sequence binding protein/IFN regulatory factor 8 drives the development of type I IFN-producing plasmacytoid dendritic cells. Journal of immunology. Feb 1 2003;170(3):1131–5. Epub 2003/01/23.

46. Kurotaki D, Kawase W, Sasaki H, Nakabayashi J, Nishiyama A, Morse HC, 3rd, et al. Epigenetic control of early dendritic cell lineage specification by the transcription factor IRF8 in mice. Blood. Apr 25 2019;133(17):1803-13. Epub 2019/02/24.

47. Dror N, Alter-Koltunoff M, Azriel A, Amariglio N, Jacob-Hirsch J, Zeligson S, et al. Identification of IRF-8 and IRF-1 target genes in activated macrophages. Mol Immunol. Jan 2007;44(4):338–46.

48. Langlais D, Barreiro LB, Gros P. The macrophage IRF8/IRF1 regulome is required for protection against infections and is associated with chronic inflammation. J Exp Med. Apr 04 2016;213(4):585–603.

49. Mancino A, Termanini A, Barozzi I, Ghisletti S, Ostuni R, Prosperini E, et al. A dual cis-regulatory code links IRF8 to constitutive and inducible gene expression in macrophages. Genes Dev. Feb 15 2015;29(4):394–408.

50. Hu X, Ivashkiv LB. Cross-regulation of signaling pathways by interferon-gamma: implications for immune responses and autoimmune diseases. Immunity. Oct 16 2009;31(4):539–50. Epub 2009/10/17.

51. Hambleton S, Salem S, Bustamante J, Bigley V, Boisson-Dupuis S, Azevedo J, et al. IRF8 mutations and human dendritic-cell immunodeficiency. The New England journal of medicine. Jul 14 2011;365(2):127–38.

52. Salem S, Langlais D, Lefebvre F, Bourque G, Bigley V, Haniffa M, et al. Functional characterization of the human dendritic cell immunodeficiency associated with the IRF8(K108E) mutation. Blood. Sep 18 2014;124(12):1894–904.

53. Bigley V, Maisuria S, Cytlak U, Jardine L, Care MA, Green K, et al. Biallelic interferon regulatory factor 8 mutation: A complex immunodeficiency syndrome with dendritic cell deficiency, monocytopenia, and immune dysregulation. J Allergy Clin Immunol. Nov 8 2017. Epub 2017/11/13.

54. Mace EM, Bigley V, Gunesch JT, Chinn IK, Angelo LS, Care MA, et al. Biallelic mutations in IRF8 impair human NK cell maturation and function. J Clin Invest. Jan 3 2017;127(1):306–20. Epub 2016/11/29.

55. Qin W, Gao J, Ma S, Wang Y, Li DM, Jiang WK, et al. Multiple Cervical Root Resorption Involving 22 Teeth: A Case with Potential Genetic Predisposition. Journal of endodontics. Oct 19 2022. Epub 2022/10/22.

56. Stossel TP, Condeelis J, Cooley L, Hartwig JH, Noegel A, Schleicher M, et al. Filamins as integrators of cell mechanics and signalling. Nat Rev Mol Cell Biol. Feb 2001;2(2):138–45. Epub 2001/03/17.

57. Razinia Z, Makela T, Ylanne J, Calderwood DA. Filamins in mechanosensing and signaling. Annu Rev Biophys. 2012;41:227–46. Epub 2012/03/13.

58. Leung R, Wang Y, Cuddy K, Sun C, Magalhaes J, Grynpas M, et al. Filamin A regulates monocyte migration through Rho small GTPases during osteoclastogenesis. J Bone Miner Res. May 2010;25(5):1077–91. Epub 2009/11/26.

59. Goldberg S, Glogauer J, Grynpas MD, Glogauer M. Deletion of filamin A in monocytes protects cortical and trabecular bone from post-menopausal changes in bone microarchitecture. Calcified tissue international. Aug 2015;97(2):113–24. Epub 2015/04/22.

60. Yang C, Yang P, Liu P, Wang H, Ke E, Li K, et al. Targeting Filamin A alleviates ovariectomy-induced bone loss in mice via the WNT/beta-catenin signaling pathway. Cell Signal. Feb 2022;90:110191. Epub 2021/11/15.

